# The mesolimbic dopamine signatures of relapse to alcohol-seeking

**DOI:** 10.1101/2020.03.06.981605

**Authors:** Yu Liu, Philip Jean-Richard-dit-Bressel, Joanna Oi-Yue Yau, Alexandra Willing, Asheeta A. Prasad, John M. Power, Simon Killcross, Colin W.G. Clifford, Gavan P. McNally

**Affiliations:** School of Psychology, UNSW Sydney, Australia; Department of Physiology and Translational Neuroscience Facility, School of Medical Sciences, UNSW Sydney, Australia

**Author notes:** Correspondence: Gavan P. McNally.

## Abstract

The mesolimbic dopamine system comprises distinct compartments supporting different functions in learning and motivation. Less well understood is how complex addiction-related behaviors emerge from activity patterns across these compartments. Here we show how different forms of relapse to alcohol-seeking are assembled from activity across the ventral tegmental area and the nucleus accumbens. Using gCaMP and dLight fibre photometry, we show that self-administration and two forms of relapse (renewal/context-induced reinstatement and reacquisition) are associated with recruitment across the mesolimbic dopamine system. Using a variety of interventions, we show that this activity is causal to both forms of relapse. Finally, we use dissimilarity matrices to identify mesolimbic dopamine signatures of self-administration, extinction, and relapse. We show that signatures of relapse can be identified from heterogeneous activity profiles across the mesolimbic dopamine system and that these signatures differ for different forms of relapse.

## Introduction

It is axiomatic that the actions of dopamine are critical to drug addiction. Dopamine mediates the reinforcing effects of drugs of abuse, instructs learning about behavioral as well as environmental antecedents to these effects, and contributes to action selection controlling drug seeking and drug taking (Everitt et al., 2008; Luscher, 2015; Nutt et al., 2015). Moreover, exposure to drugs of addiction can profoundly alter these functions (Volkow et al., 2017).

Much remains to be learned about these roles of dopamine in addiction. Chief among these is how dopamine contributes to relapse. A persistent propensity to relapse is a diagnostic feature of drug addiction and remains a primary impediment to successful long-term treatment (Association, 2013; Jonas et al., 2014). In animal models, dopamine mediates various forms of relapse to drug seeking (cue-, stress-, priming, context/renewal reinstatement) because these can be prevented by systemic or local manipulation of ventral tegmental area (VTA) and nucleus accumbens (Acb) (Bossert et al., 2013; Bossert et al., 2007; Chaudhri et al., 2009; Gibson et al., 2018; Hamlin et al., 2007; Mahler et al., 2014; McFarland and Kalivas, 2001; Schmidt et al., 2005). However, the mesolimbic dopamine system has a complex architecture. VTA dopamine neurons form distinct channels linked to differences in behavioral and motivational function (Cohen et al., 2012; de Jong et al., 2019; Heymann et al., 2019; Lammel et al., 2008; Lammel et al., 2011; Lammel et al., 2013; Saunders et al., 2018; Tian et al., 2016; Watabe-Uchida et al., 2012). In turn, there are distinct profiles of dopamine release across compartments of the ventral striatum giving rise to different functions (de Jong et al., 2019; Mohebi et al., 2019).

Relapse is presumably assembled from activity across the mesolimbic dopamine system. But how relapse relates to activity in specific VTA and accumbens compartments, whether this is the same for different forms of relapse, and how relapse-associated activity relates to activity during self-administration or extinction/abstinence are unknown. Here we answered these questions by studying the temporal dynamics of activity across spatially distinct compartments of the mesolimbic dopamine system. We show that signatures of relapse can be identified from heterogeneous activity profiles across the mesolimbic dopamine system and that these signatures differ for different forms of relapse.

## Materials and Methods

### Subjects

Subjects were adult male Long-Evans (School of Psychology, UNSW Australia) or Th-Cre (SD-Th-cre^tm1sage^) (Sage Laboratories, Cambridge, United Kingdom) rats. They were housed in ventilated racks, in groups of 4, on corn cob bedding in a climate-controlled colony room maintained on 12:12 hour light/dark cycle (0700 lights on). Rats had free access to food (Gordon’s Rat Chow) and water until two days prior to commencement of behavioral training when they received 1 hour of access to food and water each day for the remainder of the experiment. All subjects were randomly allocated to experimental conditions. All studies were performed in accordance with the Animal Research Act 1985 (NSW), under the guidelines of the National Health and Medical Research Council Code for the Care and Use of Animals for Scientific Purposes in Australia (2013). The UNSW Animal Care and Ethics Committee approved all procedures.

### Surgery

Rats were anaesthetized via i.p. injection with a mixture of 1.3 ml/kg ketamine anaesthetic (Ketamil; Troy Laboratories, NSW, Australia) at a concentration of 100 mg/ml and 0.3 ml/kg of the muscle relaxant xylazine (Xylazil; Troy Laboartories, NSW, Australia) at a concentration of 20 mg/ml. Rats received a subcutaneous injection of 0.1ml 50mg/ml carprofen (Pfizer, Inc., Tadworth, United Kingdom) before being placed in the stereotaxic frame (Kopf Instruments, CA, USA). Rats received stereotaxic surgery using the following flat skull coordinates relative to bregma (in mm):

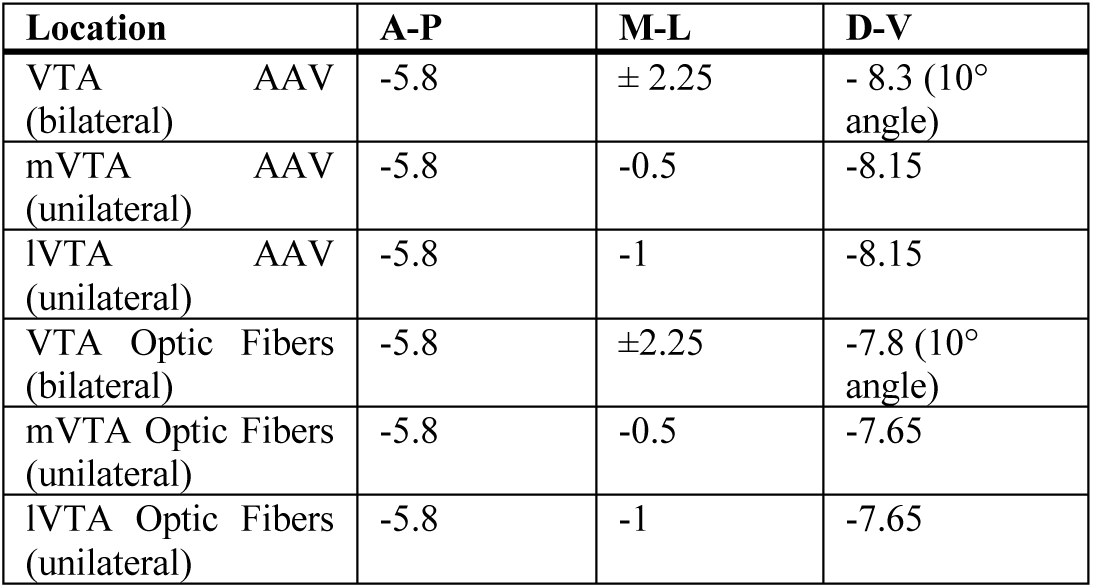
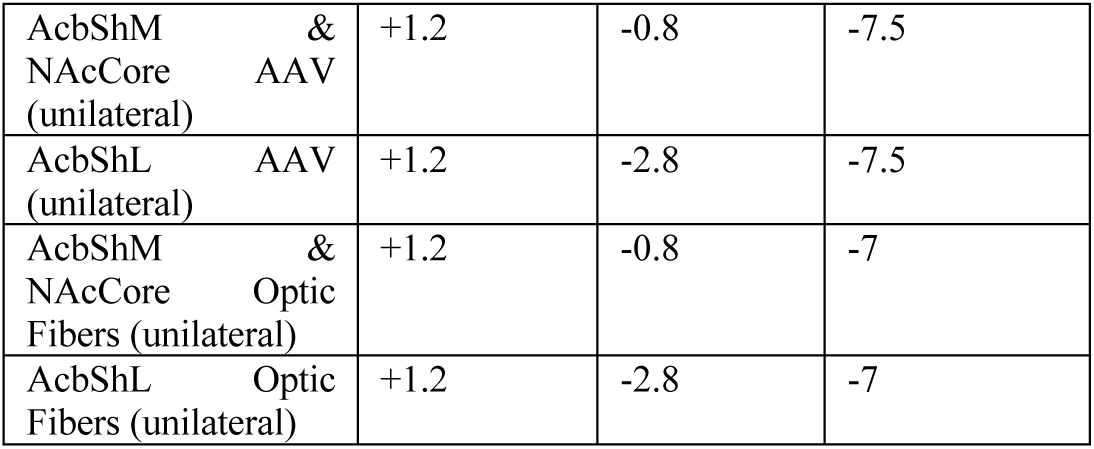

Vectors (0.5μl hM4Di and KORD, 0.75μl otherwise) were infused with a 23-gauge, cone tipped 5μl stainless steel injector (SGE Analytical Science, Australia) over 3 min using an infusion pump (UMP3 with SYS4 Micro-controller, World Precision Instruments, Inc., FL, USA). The needle was left in place for 7 min to allow for diffusion of and reduce spread up the infusion tract. Optic cannulae for relevant experiments were implanted during a concurrent stereotaxic procedure and secured using jeweller’s screws and dental cement (Vertex Dental, The Netherlands). At the end of surgery, rats received i.p. injection of 0.2 ml of 150 mg/ml solution of procaine penicillin (Benacillin; Troy Laboratories, NSW, Australia) and 0.2 ml of 100 mg/ml cephazolin sodium (AFT Pharmaceuticals, NSW, Australia). iP rats received the antibiotic baytril (3mg/kg, i.p) and the anti-inflammatory drug meloxicam (3mg/kg, i.p). All rats were monitored daily for weight and/or behavioral changes.

### Behavioural Procedures

#### Apparatus

Standard rat operant chambers (ENV-008) (Med Associates, VT, USA) with dimensions 29.5 cm (width) x 24.8 cm (length) x 18.7 cm (height) were used for all alcohol self-administration and extinction procedures. The chambers contained two nosepoke holes symmetrically located on one sidewall of the chamber, 3cm above a grid floor. A recessed magazine was located behind a 4×4cm opening in the center of the same wall between the two nosepokes. Responding on one (active) nosepoke extinguished the cue light in the nosepoke and triggered a syringe pump to deliver alcoholic beer to the magazine during acquisition training, whereas responding on the other (inactive) nosepoke had no programmed consequences. A computer running MedPC-IV software controlled all events. For optogenetic experiments, an LED plus fiber-optic rotary joint and LED driver (Doric Lenses Inc., Quebec, Canada) was suspended above each chamber and controlled by MedPC-IV. For fibre photometry, the patch cable was supported by a gimbal and counterweighted arm. The eight self-administration chambers were divided into two groups of four to serve as distinct contexts for experiments with context as a factor. These chambers differed in their olfactory (rose vs peppermint essence), tactile (grid vs Perspex flooring) and visual (light on vs off) properties. These two contexts were counterbalanced to serve as the training (context A) and extinction (context B) contexts. All fiber optic cannulae and patch cables were hand fabricated and tested using parts from Thor Labs (Newton, NJ, USA). Construction procedure was adapted from the protocol described by Sparta et al. (2012). Fiber optic cannulae and patch cables were fabricated from 0.39 NA, Ø400μm core multimode optical fiber and ceramic ferrules (Thor Labs, Newton, NJ, USA).

Locomotor chambers (ENV-515) (Med Associates, VT, USA) with dimensions 43.4 cm (width) x 43.4cm (length) x 30.3 cm (height) were used for locomotor assessment. Movement was tracked with three 16 beam infrared arrays. Infrared beams were located on both the x- and y-axes for positional tracking.

#### General behavioral testing procedures

All behavioral procedures commenced a minimum of 4 weeks after surgery. On the first two days, the rats received 20 min magazine training sessions in context A and context B, counterbalanced order. During these sessions, there were 10 non-contingent deliveries of 0.6ml of the reward (4% alcohol (v/v) decarbonated beer; Coopers Brewing Company, Australia) at time intervals variable around a mean of 1.2 min. On the next ten days rats received self-administration training in context A for 1 hour per day (unless otherwise stated). Responding on the active nosepoke extinguished the nosepoke cue light and triggered delivery via syringe pump of 0.6 ml alcoholic beer to the magazine on an FR-1 schedule followed by a 24 seconds timeout. Responses on the inactive nosepoke were recorded but had no consequences. On the next four days, rats received extinction training in context B for 1 h per day (unless otherwise stated). During this training, responses on the active nosepoke extinguished the cue light and triggered the pump but no beer was delivered.

Testing commenced 24 hours after extinction. Rats were tested for 1 hour in the extinction context (ABB) and for 1 hour in the training context (ABA) for expression of extinction and renewal (context-induced reinstatement), respectively. The order of tests was counterbalanced and tests were 24 hours apart. Tests were identical to self-administration except that the syringe pump was empty. Rats were tested 24 hours later for 1 hour reacquisition of alcoholic beer seeking in the training context. Our past research has shown no impact of the prior order of ABA and ABB testing on responding during reacquisition. Unless otherwise stated, tests lasted 60 min. We selected these procedures based on our past work that has shown robust context-induced reinstatement and reacquisition under these conditions(Hamlin et al., 2007; Gibson et al., 2018).

#### Chemogenetic hM4Di inhibition of VTA^Th^ neurons on renewal, reacquisition and locomotor activity

There were two groups of Th Cre+/- rats. Group eYFP (*n* = 6) whereas group hM4Di (*n* = 8) received AAV5-hSyn-DIO-hM4D(Gi)-mCherry bilaterally in the VTA. Rats were trained, extinguished, and tested as described above. Rats received an i.p. injection 0.1 mg/kg of clozapine (#C6305, Sigma Aldrich; 1 ml/kg diluted, 5% DMSO and saline) 15 min prior to tests. They then received two days of 30 minutes habituation each day in locomotor chambers, rats were tested for locomotor activity (30 min) 15 min after i.p. injection of saline or clozapine. The order of tests (saline or clozapine) were counterbalanced and 24 hours apart.

#### Chemogenetic KORD inhibition of VTA^Th^ neurons on reacquisition

There were two groups of Th Cre+/- rats. Group eYFP (*n* = 6) received AAV5-hSyn-DIO-eYFP whereas group hM4Di (*n* = 8) received AAV8-hSyn-DIO-KORD-mCitrine, bilaterally in the VTA. Rats were trained, extinguished, and tested as described above with the exceptions that all training and testing occurred in a single context and rats were tested for reacquisition only, 24 hours after last extinction session. Rats were injected s.c. with salvinorin B (SalB) (Apple Pharms Ingredients Inc, NC, USA; (15 mg/kg, 0.5 ml/kg) 15 min prior to reacquisition. salvinorin B was dissolved in 100% dimethyl-sulfoxide (DMSO). Rats had been habituated to the s.c. injection procedure via 4 daily injections (0.5 ml/kg, 100% DMSO).

#### Fiber photometry of Ca2+ transients in VTA^Th^ neurons during acquisition, extinction, renewal and reacquisition

There was one group of Th Cre+/- rats (*n* = 24) with fibers targeted unilaterally at the lVTA (*n* = 16) or mVTA (*n* = 8). Rats received AAV5-hSyn.Flex.GCaMP6f.WPRE.SV40 eYFP unilaterally in the VTA. Rats were trained, extinguished, and tested as described above. Fiber photometry recordings were made on Days 2 and 10 of self-administration training and Day 1 of extinction training. These sessions were 60 min duration and recordings were made for the first 30 min. Recordings were also made during tests for extinction (ABB), renewal (ABA), and reacquisition. These sessions were 30 min in duration and recordings were made for 30 min.

#### Optogenetic inhibition of VTA on renewal and reacquisition

There was one group of Th Cre-/-rats (*n* = 8) and two groups of Cre+/- rats with fibers targeted bilaterally at lVTA (*n* = 6) or mVTA (*n* = 6). Rats were trained and tested as described above, except the test sessions were 30 min in duration. During tests, rats were connected to patch cables attached to 625nm LEDs (Doric Lenses Inc., Quebec, Canada) and received optical stimulation for 10 seconds after each nosepoke during the FR (Nosepoke+) but not during the timeout (Nosepoke-). Rats had been habituated to patch cables on Days 6 and 7 of acquisition and Days 2 and 3 of extinction.

#### D1 dopamine receptor antagonist on renewal, reacquisition and locomotor activity

There 4 groups of rats injected with either 0 (*n* =8), 0.025 (*n* = 8), 0.1 (*n* = 8), or 0.25 (*n* = 8) mg/kg SCH39166. Rats were trained, extinguished, and tested as described above. Rats received s.c. injections of SCH31966 or saline 15 min prior to test sessions. Then then received two days of 30 minutes habituation to the locomotor chambers prior to being tested for locomotor activity (30min). Rats all received s.c. injection of saline and SCH39166 at the same dose they had received during relapse tests. a dose corresponding to their group. The order of these tests (saline and SCH39166) was counterbalanced and tests were 24 hours apart.

#### Fiber photometry of dopamine transients in Acb during acquisition, extinction, renewal and reacquisition

There were three groups of rats with AAV and fibers targeted unilaterally at the AcbShM (*n* = 6), AcbC (*n* = 7), or AcbShL (*n* = 5). Rats received AAV5-CAG-dLight1.1 and optic fibers unilaterally in the Acb. Fiber photometry recordings were made on Days 2 and 10 of self-administration and Day 1 of extinction training. These sessions were 60 min duration and recordings were made for the first 30 min. Recordings were also made during tests for extinction (ABB), renewal (ABA), and reacquisition. These sessions were 30 min in duration and recordings were made for 30 min.

### Fibre photometry

Recordings were performed using Fibre Photometry Systems from Doric Lenses and Tucker Davis Technologies (RZ5P, Synapse). Two excitation wavelengths, 465 nm (Ca^2+^ - dependent signal) and 405 nm (isosbestic control signal) emitted from LEDs (465nm: LEDC1-B_FC, 405nm: LEDC1-405_FC; Doric Lenses), controlled via dual channel programmable LED drivers (LEDD_4, Doric Lenses), were channeled into 0.39 NA, Ø400μm core multimode pre-bleached patch cables via a Doric Dual Fluorescence Mini Cube (FMC2, Doric Lenses). Light intensity at the tip of the patch was maintained at 10-30µW across sessions. GCaMP6, dLight1.1 and isosbestic fluorescence, wavelengths were measured using femtowatt photoreceivers (Newport, 2151). Synapse software controlled and modulated excitation lights (465nm: 209Hz, 405nm: 331 Hz), as well as demodulated and low-pass filtered (3Hz) transduced fluorescence signals in real-time via the RZ5P. Synapse/RZ5P also received Med-PC signals to record behavioral events and experimenter-controlled stimuli in real time.

### Histology and Immunohistochemistry

#### eYFP/mCherry immunohistochemistry

Rats were deeply anesthetized with sodium pentobarbital (100 mg/kg, i.p.; Virbac, NSW, Australia) and perfused transcardially with 200 ml of 0.9% saline, containing heparin (360μl/L) and sodium nitrite (12.5ml/L), followed by 400 ml of 4% paraformaldehyde in 0.1M phosphate buffer (PB), pH7.4. Brains were extracted from the skull and post fixed for one hour in the same fixative and then placed in 20% sucrose solution overnight. Brains were frozen and sectioned coronally at 40μm using a cryostat (Leica CM1950).

To visualize eYFP, mCherry. mCitrine immunoreactivity (-IR) (Rabbit anti-eGFP Polyclonal Antibody, Cat#AA11122; RRID AB_221569; Rabbit anti-mCherry Polyclonal Antibody, Cat#PA5-34974; RRID AB_2552323 ThermoFisher Scientific) four serially adjacent sets of sections from the regions of interest were obtained from each brain and stored in 0.1% sodium azide in 0.1M PBS, pH 7.2. Sections were washed in 0.1M PB, followed by 50% ethanol, 50% ethanol with 3% hydrogen peroxidase, then 5% normal horse serum (NHS) in PB (30 minutes each). Sections were then incubated in rabbit antiserum against eGFP or mCherry (1:2000; ThermoFisher Scientific, MA, USA) in a PB solution containing 2% NHS and 0.2% Triton X-10 (48 hours at 4°C). The sections were then washed and incubated in biotinylated donkey anti-rabbit (1:1000; 24 hours at 4C; Biotin Donkey Anti-Rabbit Cat#711-065-152; RRID:AB_2540016 Jackson ImmunoResearch Laboratories, PA, USA). Finally, sections were incubated in avidin-biotinylated horseradish peroxidase complex (6 μl/ml avidin and 6 μl/ml biotin; 2 hours at room temperature; Vector Laboratories, CA, USA), washed in PB, and then incubated for 15 minutes in a diaminobenzidine solution (DAB) containing 0.1% 3,3-diaminobenzidine, 0.8% D-glucose and 0.016% ammonium chloride. Immunoreactivity was catalyzed by the addition of 0.2μl/ml glucose oxidase aspergillus (24 mg/ml, 307 U/mg, Sigma-Aldrich, NSW, Australia). Brain sections were then washed in PB, mounted onto gelatin-coated slides, dehydrated, cleared in histolene, and coverslipped with Entellan (Proscitech, Kirwin, Australia), and assessed using an Olympus BX50 transmitted light microscope (Olympus, Shinjuku, Tokyo, Japan) or Axio Scan.Z1 slide scanner (Zeiss, Oberkochen, Germany).

Criteria for inclusion in final analysis was correct AAV and/or fiber placements determined after histology.

#### Double fluorescence immunohistochemistry

Rats were deeply anesthetized with sodium pentobarbital (100 mg/kg, i.p.; Virbac, NSW, Australia) and perfused transcardially with 200 ml of 0.9% saline, containing heparin (360μl/L) and sodium nitrite (12.5ml/L), followed by 400 ml of 4% paraformaldehyde in 0.1M phosphate buffer (PB), pH7.4. Brains were extracted from the skull and post fixed for one hour in the same fixative and then placed in 20% sucrose solution overnight. Brains were frozen and sectioned coronally at 40μm using a cryostat (Leica CM1950). Four serially adjacent sets of sections from the regions of interest were obtained from each brain and stored in 0.1% sodium azide in 0.1M PBS, pH 7.2. Free-floating sections were washed repeatedly in 0.1M PBS (pH 7.2), followed by a 2h incubation in PBS (pH7.2) containing 10% normal donkey serum (NDS) and 0.2% Triton X-100. Sections were then incubated in the primary antibodies diluted in 0.1M PBS (pH 7.2) containing 0.1% sodium azide, 2% NHS and 0.2% Triton X-100, for 48h at room temperature, with gentle agitation. The primary antibodies used were sheep anti TH (1:1000; Sheep Anti-Tyrosine Hydroxylase Polyclonal Antibody; Cat # PA1-4679;

RRID: AB_561880; ThemoFisher Scientific, MA, USA) and rabbit anti-eGFP (1:1000; Rabbit anti eGFP Polyclonal Antibody; Cat#A-11122; RRID: AB_221569; ThemoFisher Scientific, MA, USA). After washing off unbound primary antibodies, sections were then incubated for 4hr at room temperature in secondary antibodies diluted in 0.1 M PBS (pH 7.2) containing 2% NHS and 0.2% Triton X-100 (PBST-X).). The secondary antibodies used were Alexa 488 donkey anti rabbit (1:500; Donkey anti-Rabbit IgG (H+L) Highly Cross-Adsorbed Secondary Antibody, Alexa Fluor 488; Cat# A-21206; RRID: AB_2535792; ThemoFisher Scientific, MA, USA) and Alexa 594 donkey anti goat (1:500; Donkey anti-Goat IgG (H+L) Cross-Adsorbed Secondary Antibody, Alexa Fluor 594; Cat# A-11058; RRID: AB_2534105; ThemoFisher Scientific, MA, USA). After washing off unbound secondary antibodies sections were mounted onto gelatin-treated slides and coverslipped with Permafluor mounting medium (Thermofisher Scientific, MA, USA). Fluorescent images were taken by Olympus BX53 upright microscope (Olympus, Shinjuku, Tokyo, Japan).

### Statistical Analyses

#### Inclusion criteria

The criteria for inclusion in final analyzes were correct AAV and fiber placements as determined after histology, so animals were excluded if either AAV or fiber optic cannulae were misplaced. Data in figures are represented as mean ± SEM unless otherwise stated. Group sizes were based on past experience with these preparations showing that they were sufficient to detect large (d = 0.8) effect sizes in behavioral studies with at least 80% power. Group numbers used for analyzes in each experiment are indicated at two locations: (1) under the subheadings of behavioral procedures above; (2) in the main results text.

#### Behavior

Our primary behavioral dependent variables were numbers of active nosepokes, inactive nosepokes, and distance travelled (locomotor activity). These were analyzed by means of ANOVA and analyzes involving repeated measures adopted a multivariate approach (Harris, 2004). All analyzes partitioned variances into main effect and interaction terms using Psy Statistical Package (Bird, 2004).

#### Fiber Photometry

For fiber photometry, the primary dependent variables were Ca2+ or dopamine transients around nosepokes and magazine entries during training (sessions 2 and 10), extinction (session 1), extinction test (ABB), renewal test (ABA), and reacquisition. These data were analyzed using custom MATLAB scripts.

Specifically, for gCaMP experiments, Ca2+ dependent and Ca2+ independent (isosbestic) signals during recording sessions were extracted and downsampled (15.89 samples/sec); signals around logged disconnections were removed prior to further signal processing. The isosbestic signal was regressed onto the Ca2+ -dependent signal to create a fitted isosbestic signal and a fractional fluorescence signal ΔF/F was calculated via subtracting fitted 405 nm signal from 465 nm channels, and then dividing by the fitted 405 nm signal. This produces a motion-artifact-corrected Ca2+ signal with a mean of ∼0. Signal-to-noise was further boosted via filtering after contributions of low and high frequency components were determined via fast fourier transform; ΔF/F signal was convolved over 90sec (high-pass filtered) and then low-pass filtered at 2Hz.

ΔF/F within a time window around events was compiled and averaged for each trials To determine significant event-related transients within this window, a bootstrapping confidence interval (CI) procedure (95% CI, 1000 bootstraps) was used (Jean-Richard-dit-Bressel et al., 2020). A distribution of bootstrapped ΔF/F means was generated by randomly resampling from trial ΔF/F waveforms, with replacement, for the same number of trials. A confidence interval was obtained per timepoint using the 2.5 and 97.5 percentiles of the bootstrap distribution, which was then expanded by a factor of sqrt(n/(n-1)) to adjust for narrowness bias. Significant transients were defined as periods whose 95% CI did not contain 0 continuously (minimum 1/2sec).

A similar analysis procedure was used for dLight experiments, signals (465nm, 405nm) were downsampled and processed (logged disconnections removed, ΔF/F obtained using fitted isobestic). Then, signals were low-pass filtered at 3Hz (significant transients minimum = 1/3 sec) instead of 2Hz, due to the faster dynamics of dLight1.1 compared to gCaMP.

Waveform kernels were obtained by normalising each trial waveform according to its sum square deviation from 0 and were used for further analysis. This approach to normalisation preserves the baseline of the waveform and allows interpretation of the mean kernel when it is different from 0.

#### Representational similarity analysis

Representational similarity analysis (RSA; Kriegeskorte and Bandettini, 2008) is a modality-independent way of comparing activity patterns. To this end, the dissimilarity of two peri-event kernel waveforms was quantified using correlation distance (1 – Pearson correlation) of the mean trial kernel waveforms (±3s around events). A first order representational dissimilarity matrix (RDMs) was formed from the pairwise correlation distances, indicating the degree to which each pair of waveforms are similar/dissimilar. First order RDMs representing dissimilarity in activity patterns to Nosepoke+ across session for each brain region (Figure 2D,2E, Figure 3D, 3E, 3F) were generated using custom MATLAB scripts. A cross-experiment first order RDM was constructed using the same methodology (correlation distance of peri-event activity), using the mean Nosepoke+ kernel across sessions and brain regions as inputs. The dissimilarity in the dissimilarity matrices were assessed using second order RSA (Kriegeskorte and Bandettini, 2008; Figure 4). Instead of comparing neural activity, second order RSA compares RDMs. By abstracting from underlying data, second order RSA is modality independent and can be used to compare similarities in disparate data types (e.g. fMRI, neuron spiking). For dissimilarity of different brain region activity profiles (Figure 4B), the correlation distance (1 – Spearman correlation) between single brain region RDMs (cross-session RDM) was calculated. For dissimilarity of different session activity profiles (Figure 4C), the correlation distance (1 – Spearman correlation) between single session RDMs (cross-region RDM) was calculated.

**Figure 1.**
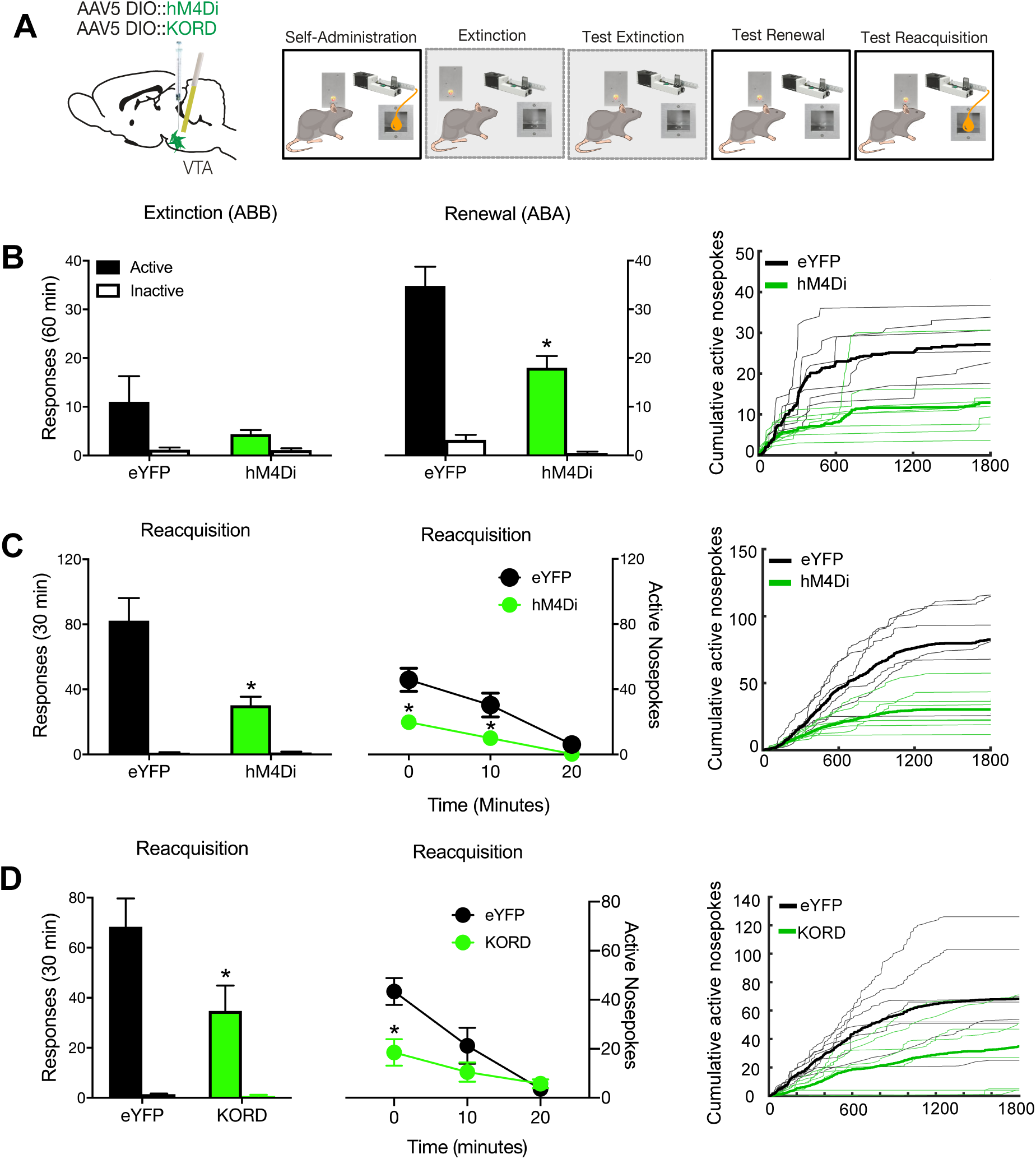
VTA TH neurons mediate relapse. (**A**) Cre-dependent inhibitory DREADDs were applied to the VTA. Rats were trained and tested in an ABA renewal procedure and also tested for reacquisition. (**B**) hM4Di chemogenetic inhibition of VTA TH neurons reduced responding during ABA renewal (context-induced reinstatement) of alcohol seeking. (**C**) hM4Di chemogenetic inhibition of VTA TH neurons reduced reacquisition, with data shown as total and in 10 min time bins. (**D**) KORD chemogenetic inhibition of VTA TH neurons also reduced reacquisition, with data shown as total and in 10 min time bins. Raw data are shown for all individual rats as cumulative functions. All analyses were ANOVAs. * p < .05.

**Figure 2.**
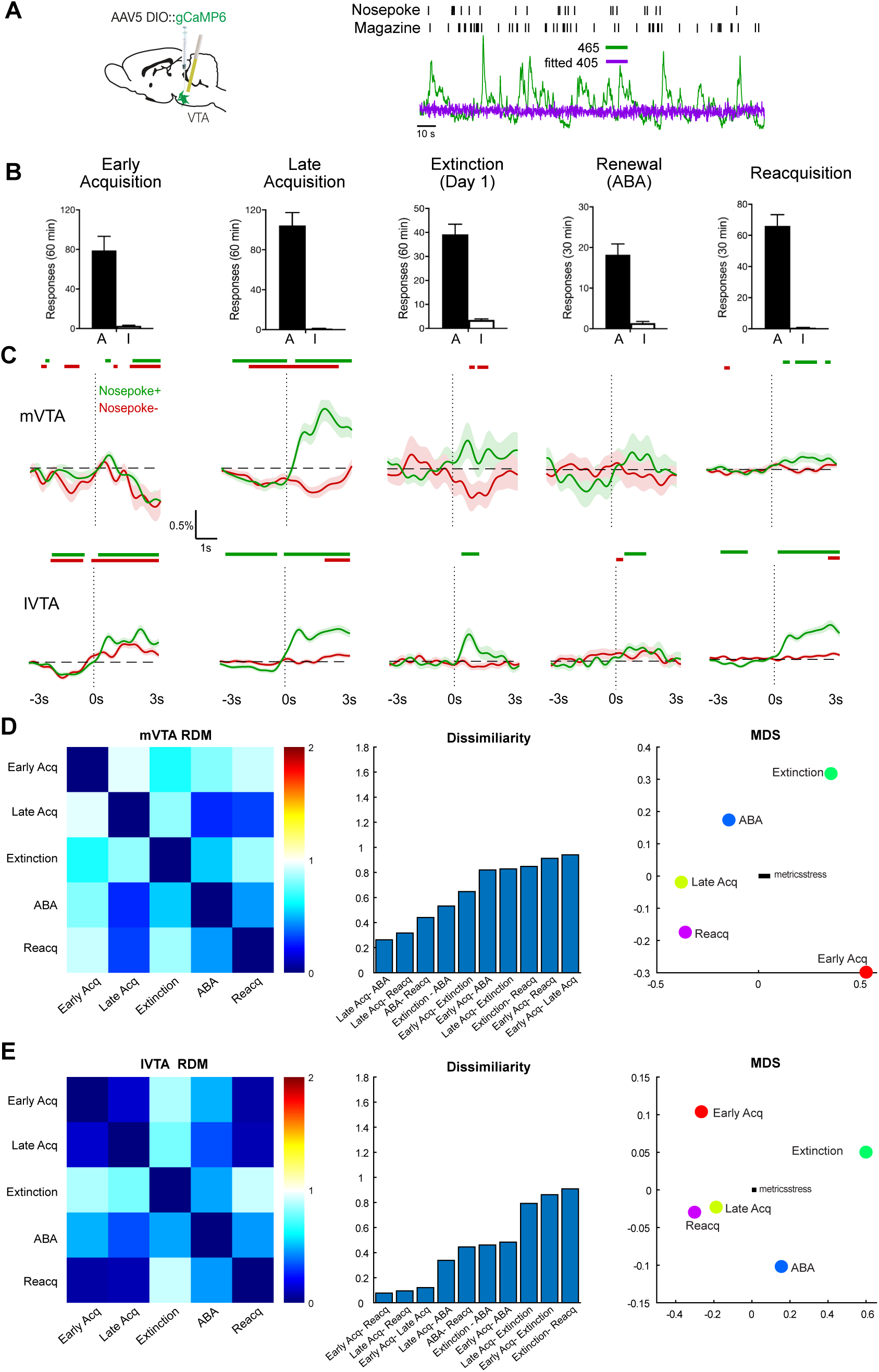
Activity of VTA TH neurons during relapse. (**A**) Cre-dependent gCaMP6f was applied to the VTA and rats were trained and tested in an ABA renewal procedure and for reacquisition. A representative trace from recording is shown. (**B**) Mean ± SEM active, *A*, and inactive, *I*, nosepokes from the five recording sessions. (**C**) Mean ± SEM calcium transients in VTA TH neurons ±3 s around Nosepoke+ (active nosepokes that triggered the pump) and Nosepoke-(active nosepokes at other times) for mVTA and lVTA. Coloured bars above traces show periods (with minimum consecutive threshold) of significant difference from 0 as defined by 95% confidence intervals. (**D**) and (**E**), Representational dissimilarity matrix (RDM) for mVTA and lVTA (0 for perfect similarity, 1 for no similarity, 2 for perfect dissimilarity) with multidimensional scaling (MDS) visualisation whereby distances reflect dissimilarity.

**Figure 3.**
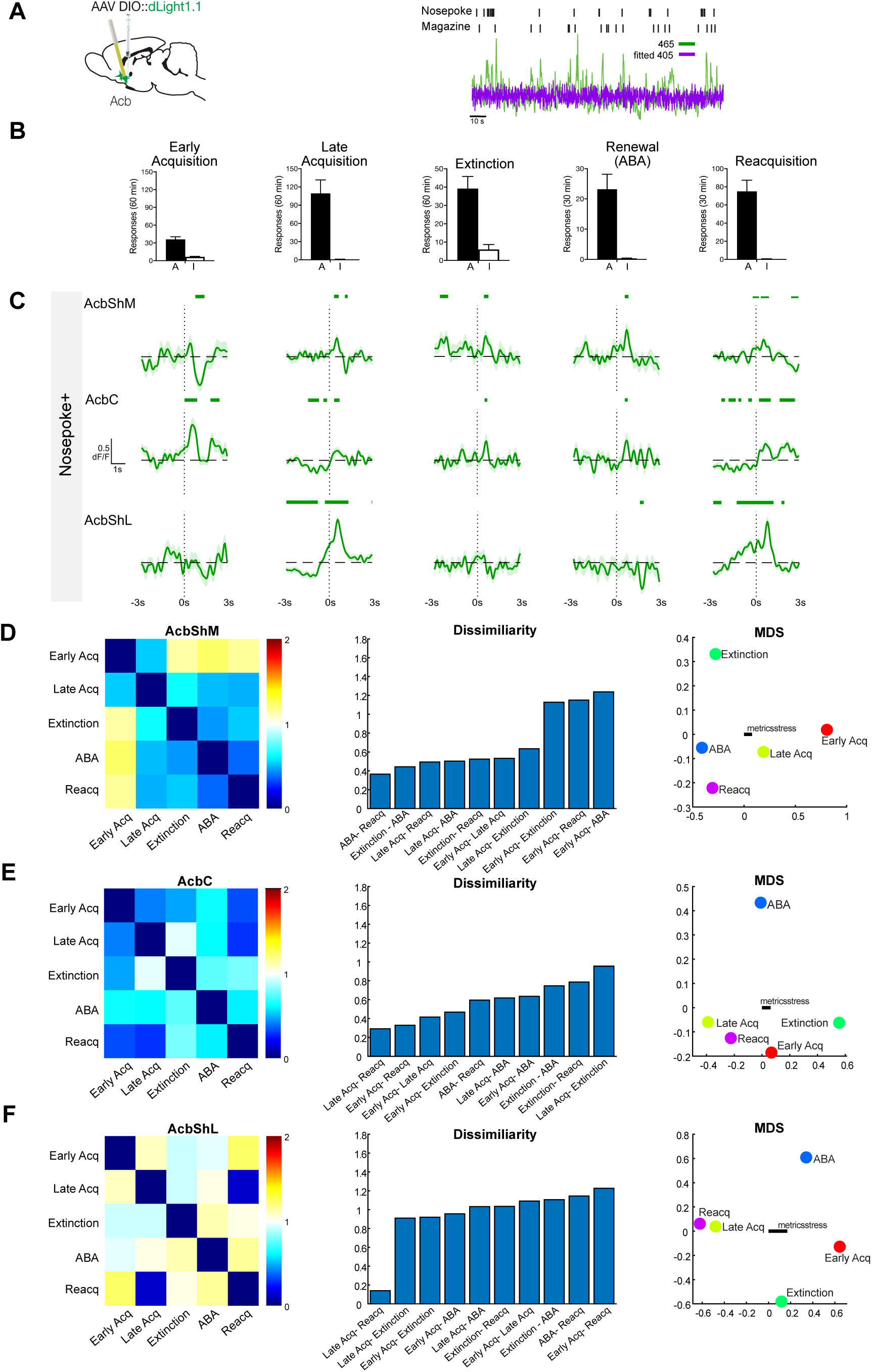
Dopamine transients in the ventral striatum. **A**) dLight was applied to AcbShM, AcbC, or AcbShL and rats were trained and tested in an ABA renewal procedure and for reacquisition. A representative recording trace is shown. (**B**) Mean ± SEM active, *A*, and inactive, *I*, nosepokes from the five recording sessions. (**C**) Mean ± SEM dopamine transients ±3 s around Nosepoke+. Green bars above traces show periods (with minimum consecutive threshold) of significant difference from 0 as defined by 95% confidence intervals. (**D** - **F**), RDMs with multidimensional scaling (MDS) visualisation whereby distances reflect dissimilarity.

**Figure 4.**
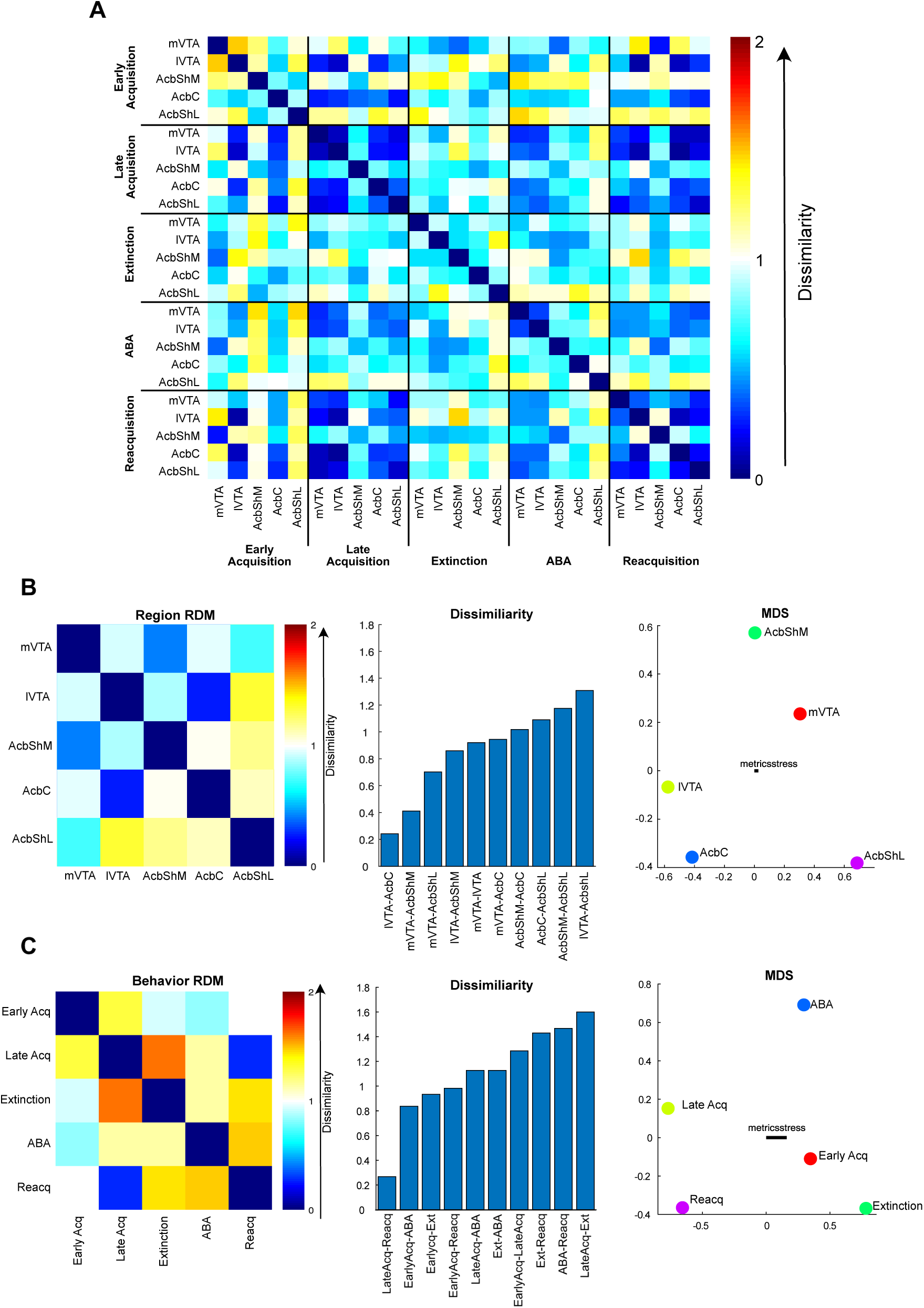
Representational similarity analysis. (**A**), First-order RDM showing correlation distances between all pairs of conditions. (**B**) Second order Brain RDM showing dissimilarity for each pair of brain regions, and MDS visualisation of these differences whereby distances reflect dissimilarity. **(C**) Second order Behavior RDM showing dissimilarity for each pair of behavioral self-administration stages, difference from RDM for each pair of behavioral conditions, and MDS visualisation of these differences.

To visualise first and second order similarities/dissimilarities between pairs of condition, correlation distances were also conveyed using multidimensional scaling. Coordinates were obtained using (MATLAB mdscale function (p = 2, criterion = metricsstress). Across graphs, S-Stress was consistently below 0.2, indicating a fair fit of the data.

## Results

### Ventral tegmental area TH neurons mediate different forms of relapse

First, we studied the causal roles of VTA^TH^ neurons in relapse. To do this, we expressed the cre-dependent inhibitory hM4Di DREADD) (*n* = 8) or enhanced yellow fluorescent protein (eYFP) (*n* = 6) in the VTA of TH-Cre rats (Liu et al., 2016) (**Figure S1A, C**), then trained rats to respond for alcoholic beer in a distinctive context (A). Responses on one nosepoke (active) caused delivery of alcoholic beer to a magazine cup whereas responses on a second nosepoke (inactive) did not. In each experiment, rats readily learned to self-administer (**Table S1**). Then we extinguished this behavior in a second, distinctive context (B) where responses did not earn alcohol. In each experiment, responding was reduced by extinction training (**Table S1**).

We tested for two forms of relapse: renewal (context-induced reinstatement) (Crombag and Shaham, 2002; Hamlin et al., 2007; Bouton and Todd, 2014) and reacquisition (Willcocks and McNally, 2011). For renewal, rats were tested in the extinction context (extinction, ABB test) and in the training context (ABA) (**Figure 1A**). There was no alcohol available during these tests and rats were injected with the hM4Di ligand clozapine (0.1 mg/kg, i.p.). Rats refrained from alcohol-seeking in the extinction context but relapsed to alcohol-seeking in the training context (context main effect: F (1, 12) = 52.91, p < 0.001; nosepoke main effect F (1, 12) = 72.77, p < 0.001; context x nosepoke interaction, F (1, 12) = 39.97, p < 0.001) (**Figure 1B**), showing renewal (context-induced reinstatement) of alcohol-seeking. VTA TH chemogenetic inhibition reduced responding on ABA test (main effect group F (1, 12) = 13.01, p = 0.004; group x context interaction F (1, 12) = 5.78, p = 0.033; group x nosepoke F (1, 12) = 8.09, p = 0.015 interaction) but had no effect on locomotor function (**Figure S1D**).

Next we re-trained rats to self-administer alcohol in a single session to model the relapse provoked by contingent re-exposure to alcohol. In humans this causes a rapid return to pre-abstinence levels of drinking that is resistant to pharmacotherapy (Marlatt and Donovan, 2005). In rodents, a similar return to pre-extinction levels of self-administration is observed (Willcocks and McNally, 2011) (**Figure 1C**) (nosepoke main effect: F (1, 12) = 82.62, p < 0.001) and this was reduced by chemogenetic VTA TH inhibition (main effect group F (1, 12) = 16.09, p = 0.002; group x nosepoke interaction F (1, 12) = 15.99, p = 0.002).

In order to exclude the possibility that clozapine as the hM4Di ligand (Gomez et al., 2017) interacted with the effects of alcoholic beer during reacquisition, we expressed the cre-dependent inhibitory KORD DREADD (Marchant et al., 2016; Vardy et al., 2015) (*n* = 8) or enhanced yellow fluorescent protein (eYFP) (*n* = 6) in the VTA of TH-Cre rats and inhibited TH neurons via injection of SalB (15 ml/kg, i.p.). KORD chemogenetic inhibition of VTA TH neurons also reduced relapse during reacquisition (main effect group F (1, 14) = 5.26, p = 0.038; group x nosepoke interaction F (1, 14) = 4.55 p = 0.051 overall; F (1, 14) = 9.647, p = 0.008 first 10 min). (**Figure 1D**).

### Activity of VTA TH neurons during self-administration and relapse

Next we used fibre photometry (Gunaydin et al., 2014) to determine when during self-administration and relapse VTA TH neuron activity was important and whether this varied across the medial-lateral extent of the VTA (Lammel et al., 2011; Lammel et al., 2013; Yang et al., 2018) (**Figure 2A**). We expressed cre-dependent gCaMP6f and implanted optical fibres above VTA of TH-cre rats (n= 24). Using the lateral edge of the fasciculus retroflexus as a boundary (Ikemoto, 2007), histology showed that cannulae were located above medial (n = 8) or lateral (n = 16) VTA (**Figure S2A**). Ca^2+^ transients were recorded during early and late self-administration, the first day of extinction training, and tests (ABA renewal, ABB extinction, reacquisition) (**Figure 2B**).

We used a bootstrap 95% confidence interval procedure (bCI) (Jean-Richard-dit-Bressel et al., 2020) to analyse perievent (±3 s) df/f waveforms and detected significant Ca^2+^ transients associated with nosepokes and magazine entry. Transients were greater for active nosepokes that turned on the pump causing delivery of alcoholic beer (Nosepoke+) (**Figure 2C**) and magazine entries after a Nosepoke+ (**Figure S2B**) compared to active nosepokes or magazine entries at other times (Nosepoke- and Magazine-). Transients increased across the course of self-administration training, reduced during extinction, and were restored during renewal and reacquisition. There were obvious differences between mVTA and lVTA. There were negative transients in mVTA during early acquisition but positive transients in lVTA. By late acquisition there were positive transients in both regions. There were transients in lVTA, but not mVTA, during extinction and renewal. There were transients in both regions during reacquisition. Similar findings were observed for Magazine+ behavior **(Figure S2B**).

These findings show that relapse is associated with recruitment of VTA TH neurons. They suggest that different forms of relapse might be dissociable within VTA, with renewal linked to lVTA and reacquisition to both mVTA and lVTA. To test this, we expressed Cre-dependent eNpHR3.0 in VTA of TH Cre+ (n = 12) or Cre- (n = 8) rats and targeted optical fibres towards either medial (n = 6 TH-Cre +) or lateral (n = 6 TH-Cre+) VTA to limit optogenetic inhibition to these regions (**Figure S3A**). We silenced VTA TH neurons for 10 s only during Nosepoke+ behavior to inhibit these neurons during the time we had observed significant Ca^2+^ transients. Consistent with our photometry results, mVTA inhibition reduced reacquisition whereas lVTA inhibition reduced reacquisition and renewal (**Figure S3B**).

Event-related analyses link VTA TH activity to discrete alcohol-seeking and relapse behaviors. Indeed, these identified different roles for mVTA and lVTA that we confirmed via optogenetic inhibition. However, these analyses do not show whether or how VTA TH activity changes across self-administration, extinction, and relapse. To answer this, we compared VTA gCaMP activity patterns across different stages of the experiment using Representational Similarity Analyses (RSA) (Kriegeskorte and Kievit, 2013; Kriegeskorte et al., 2008a; Kriegeskorte et al., 2008b; Nili et al., 2014). We computed normalized Ca^2+^ waveforms for each stage (**Figure S2C**), correlated these across each pair of conditions, and computed correlation distance (1 – Pearson correlation) between pairs to measure their dissimilarity (0 for perfect correlation, 1 for no correlation, 2 for perfect anticorrelation). This complements single event analysis because it assesses dissimilarity between VTA TH activity across the experiment. Correlation distances were assembled into a first-order dissimilarity matrix (RDM) reporting dissimilarity of mVTA and lVTA TH activity signatures across the experiment (**Figure 2D, E**). Multidimensional scaling visualized these dissimilarities. There was pronounced change in the mVTA TH response across self-administration because mVTA activity in early and late self-administration was highly dissimilar. Moreover, in mVTA, both forms of relapse were more similar to established than early self-administration. For lVTA, there was less change with acquisition and different forms of relapse similar to each other.

### Dopamine release in the ventral striatum during self-administration and relapse

D1 dopamine receptors (Crombag et al., 2002), especially those in the nucleus accumbens (Acb) (Bossert et al., 2007; Chaudhri et al., 2009; Hamlin et al., 2007), are important for relapse and we confirmed this (**Figure S4A-C**). Single unit (Carelli and Deadwyler, 1994; Carelli and Ijames, 2000; Chang et al., 1998; Chang et al., 2000; Janak et al., 1999; Woodward et al., 2000) and fast-scan cyclic voltammetry (Phillips et al., 2003; Stuber et al., 2005a; Stuber et al., 2005b) recordings strongly link activity in the nucleus accumbens core (AcbC) to drug seeking, but the spatiotemporal profiles of dopamine release across the Acb during relapse are unknown as is whether these differ across forms of relapse. We used dLight photometry (Patriarchi et al., 2018) to address this (**Figure 3A**). We expressed dLight1.1 in Acb (N = 18) and implanted optical fibers above medial accumbens shell (AcbShM, n = 6), AcbC, n = 7, or lateral accumbens shell (AcbShL) (n = 5) of Long-Evans rats (**Figure S4D**). Dopamine transients were recorded during early and late self-administration, early extinction training, and relapse (**Figure 3B**).

95% bCI analyses of normalized peri event (±3 s) dLight waveforms detected significant dopamine transients for Nosepoke+ and Magazine+ (**Figure 3C, Figure S4E**) that depended on Acb subregion and the stage of the experiment. For AcbShM, there were reductions in dopamine transients early in acquisition replaced by positive transients in late self-administration and these were preserved during the remaining stages. This is similar to the profile observed for mVTA gCaMP. For AcbC, there were positive dopamine transients across each stage, similar to the profile observed for lVTA gCaMP. For AcbShL, positive transients emerged across self-administration, were lost across extinction, and restored only during reacquisition.

First-order RDMs and multidimensional scaling captured dissimilarity of Acb dopamine responses across sessions. There was pronounced change in the AcbShM dopamine response because early self-administration was dissimilar to all other stages except late self-administration. (**Figure 3D**). There was strong similarity between AcbC dopamine responses across all stages except extinction (**Figure 3E**). In marked contrast, AcbShL dopamine responses were highly dissimilar across the experiment, with only late self-administration and reacquisition similar (**Figure 3F**).

### Identifying and comparing mesolimbic dopamine signatures

Our results show recruitment across the mesolimbic dopamine system, with distinct profiles of activity within and across distinct compartments of this system. These findings raise fundamental questions about how activity in these distinct compartments relate to each other and how relapse is assembled from these distinct spatiotemporal activity profiles.

We used RSA on our complete photometry dataset to answer these questions. RSA quantitatively assesses the extent to which the dopamine activity signatures (i.e. VTA TH gCaMP and Acb dLight) across brain regions or behavioral stages are alike (Kriegeskorte and Kievit, 2013; Kriegeskorte et al., 2008b; Nili et al., 2014). We computed a first-order RDM for each brain region from early self-administration to relapse (**Figure 4A**). This reports dissimilarity between pairs of activity patterns. Then, we computed a second-order RDM across the first-order RDM for each brain region (**Figure 4B**), reporting dissimilarity between pairs of values in the first-order RDM (i.e. dissimilarity between brain regions), where dissimilarity is defined as the correlation distance (1 - Spearman correlation). Because the second-order RDMs compare first-order RDMs, it overcomes the problem of correspondency between spatiotemporally distinct activity profiles. Again, to aid interpretation, we used multidimensional scaling to visualize dissimilarity of these second-order RDMs.

Consistent with known connectivity (de Jong et al., 2019; Lammel et al., 2011; Lammel et al., 2013; Yang et al., 2018), the Region RDM (**Figure 4B**) showed that across self-administration to relapse, the signatures of lVTA and AcbC as well as mVTA and AcbShM were highly similar. Surprisingly, the signatures of the three Acb regions were highly dissimilar. In fact, there was more similarity between individual VTA and Acb subregions than among the Acb subregions themselves.

Finally, we computed second-order RDMs across experimental stage to identify and compare mesolimbic dopamine signatures for self-administration stages. The second order Behavior RDM (**Figure 4C**) identified four key findings. First, there was a change in the mesolimbic dopamine signature from early to late self-administration. Second, there were distinct mesolimbic dopamine signatures for the two forms of relapse. Third, the signature of reacquisition was similar to the signature of late self-administration. Fourth, there was little meaningful relationship between mesolimbic dopamine signatures when animals could only use environmental cues to guide their behavior because the outcome was absent (i.e. Extinction and ABA).

## Discussion

It is well established that there are heterogenous response profiles across the mesolimbic dopamine system that serve distinct functions (Cohen et al., 2012; de Jong et al., 2019; Heymann et al., 2019; Lammel et al., 2008; Lammel et al., 2011; Lammel et al., 2013; Mohebi et al., 2019; Saunders et al., 2018; Tian et al., 2016; Watabe-Uchida et al., 2012). Our findings support this. We show considerable diversity in spatiotemporal activity profiles across the VTA and ventral striatum during self-administration and two forms of relapse to alcohol-seeking. Moreover, we show that this activity causes relapse during renewal and reacquisition.

Although the isolated response profiles of individual compartments of the mesolimbic dopamine system are interesting (e.g., Figures 2C, 3C), the challenge is to move beyond individual profiles to understand how they orchestrate complex behaviors. This has proved difficult because it requires comparison of response profiles across different brain regions and different measures, in the same or different animals. This problem is not solved by multisite recording in the same animals because these still require comparison of response profiles across different brain regions and/or different sensors. RSA overcomes this problem to reveal how different features of alcohol-seeking relate to different features of the mesolimbic dopamine response. It allowed us to summarise mesolimbic dopamine signatures of relapse.

Established alcohol self-administration was a period of striking conformity across the mesolimbic system. This conformity was not present during early self-administration and instead emerged across training. Extinction, on the other hand, was a fracture point in these signatures. The similarity in activity profiles that was so pronounced during late self-administration was lost during extinction, revealing functional segregation across the mesolimbic dopamine system. Overall, there was strong similarity between mVTA and AcbShM as well as between lVTA and AcbC, consistent with contemporary understanding of mesolimbic dopamine architecture (de Jong et al., 2019; Lammel et al., 2011; Lammel et al., 2013; Yang et al., 2018). The activity signature of AcbShL, on the other hand, was highly dissimilar to the rest of the ventral striatum. This aligns with anatomical (Ikemoto, 2007), functional (Basso and Kelley, 1999; de Jong et al., 2019; Ikemoto et al., 2005), and cellular (Meredith et al., 1993) differences between AcbSh subregions, although it is worth noting that dopamine actions in both AcbShM and AcbShL are necessary for renewal of opiate seeking (Bossert et al., 2007).

Different forms of relapse have distinct mesolimbic dopamine signatures. This stands in marked contrast to influential theoretical (Bouton, 2002; Bouton, 2014; Bouton and Todd, 2014; Todd et al., 2014) and clinical (Marlatt, 1996; Stout et al., 1996) models. The mesolimbic dopamine signature for renewal was unique. It was dissimilar to self-administration, extinction, and reacquisition. This was surprising because renewal shares key behavioral features with each of these other stages. For example, renewal involves return to the self-administration context, shares with extinction the use of environmental cues to guide behavior, and shares with reacquisition a return to responding after extinction. It is precisely the similarities between these different forms of relapse that have shaped contemporary theoretical and clinical understanding. Nonetheless, the mesolimbic dopamine signature of renewal was dissimilar to these and other stages of self-administration. Relapse initiated by contextual cues is distinctly represented by the mesolimbic dopamine system.

Reacquisition, on the other hand, was not distinctly represented. The mesolimbic dopamine signature of reacquisition was remarkably similar to late self-administration, showing that extinction did not return the mesolimbic dopamine system to a naive state. Rather, the mesolimbic dopamine signature that emerged across self-administration rapidly re-appeared during reacquisition. This preservation or savings of the mesolimbic dopamine signature from late self-administration offers a powerful explanation for why relapse after contingent contact with alcohol is so difficult to prevent or treat (Anton et al., 2006; Marlatt and Donovan, 2005; Willcocks and McNally, 2014).

It remains to be discovered whether the dopamine signatures of other forms of relapse (cue-, stress-, priming) have the same or different features to those discovered here. Likewise, it remains to be discovered whether and how these signatures vary across different drugs of abuse. The RSA approach offers a powerful and straightforward way to answer these and related questions. Regardless, our findings that there are distinct mesolimbic dopamine signatures for different forms of relapse highlight the need for new theoretical models to better understand the mechanisms for relapse and to inform clinical approaches for preventing relapse.

---

**Figure S1.**
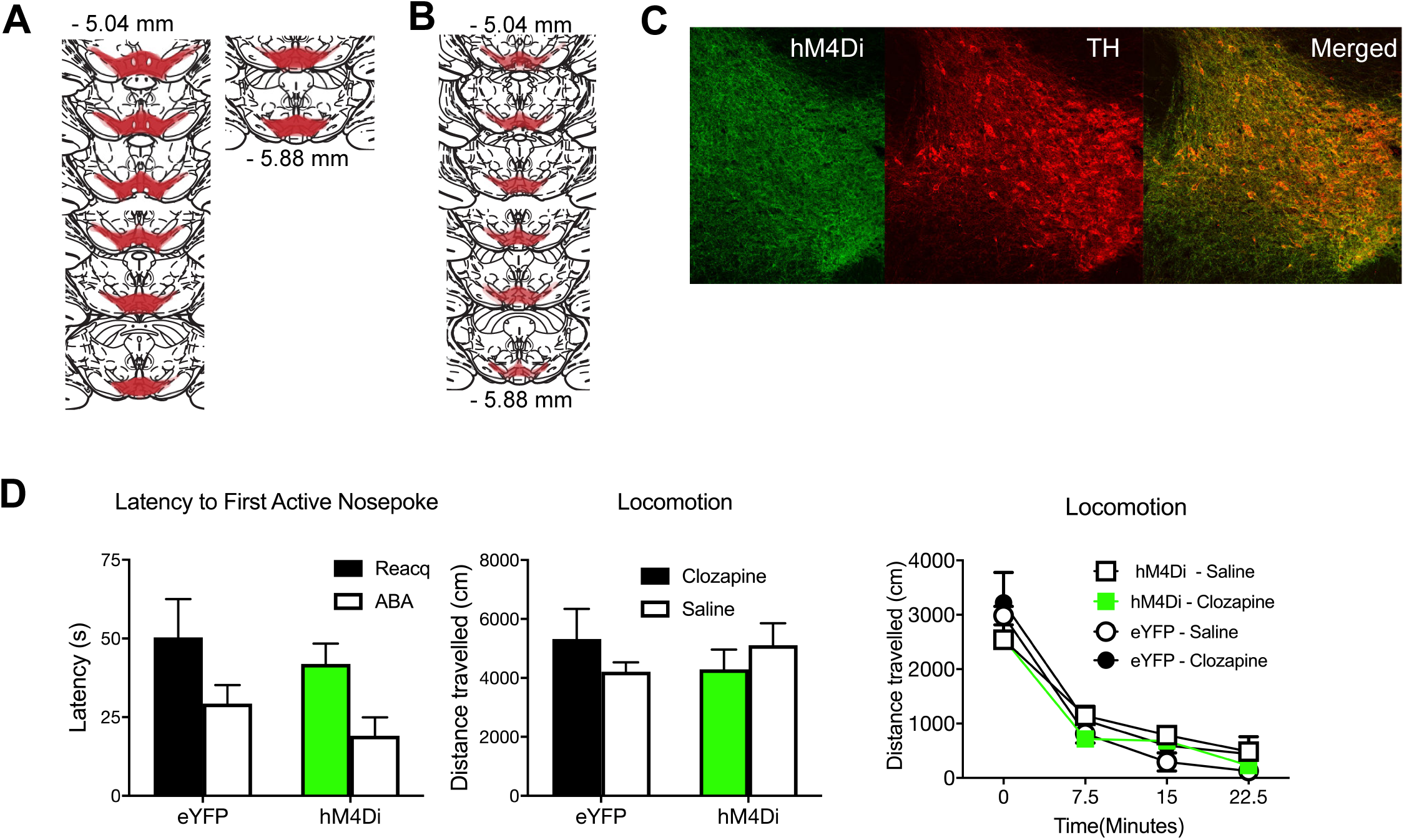
VTA TH neurons mediate relapse. (**A**) Location of hM4Di expression in midbrain with each rat represented at 25% opacity. (**B**) Location of KORD expression in midbrain with each rat represented at 25% opacity. (**C**) Example of hM4Di expression in VTA TH neurons. (**D**) No effects of hM4Di VTA TH neuron inhibition on latency to first active nosepoke during renewal or reacquisition or on locomotor activity.

**Figure S2.**
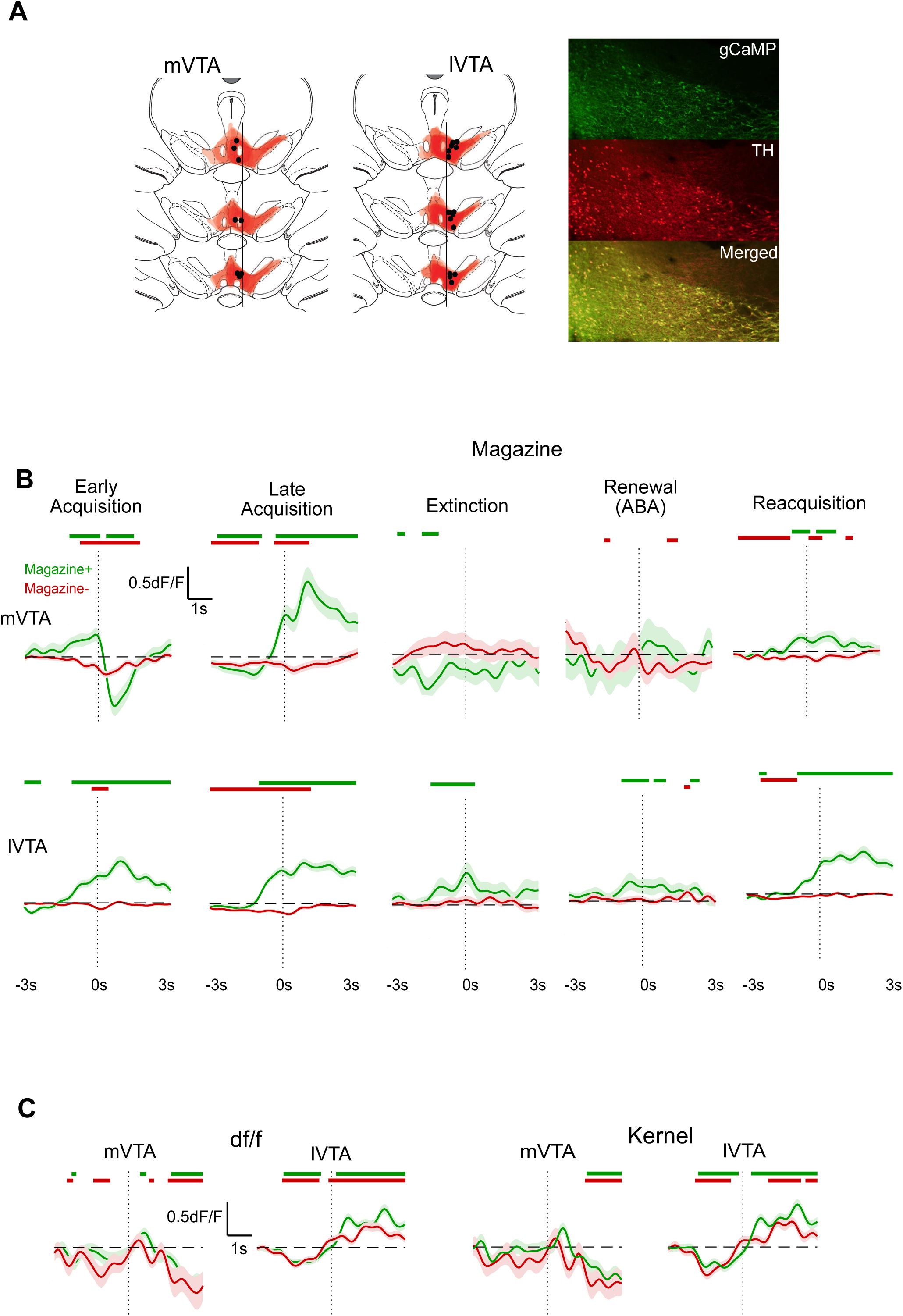
VTA fibre photometry. (**A**) Location of gCaMP6 expression and fibre tips in midbrain with each rat represented at 25% opacity. The lateral edge of the fascicuus retroflexus was used as anatomical boundary between mVTA and lVTA. Example of gCaMP6 expression in VTA TH neurons. (**B**) Mean ± SEM calcium transients in VTA TH neurons from -3s to 3 s around Magazine+ (first magazine entry after Nosepoke+) and Magazine- (magazine entries at other times) for mVTA and lVTA. Coloured bars above traces show periods (with minimum consecutive threshold) of significant difference from 0 as defined by 95% confidence intervals. (**C**) Comparison of actual df/f and normalized waveform kernels. Normalized kernels were obtained by normalising each trial waveform according to its sum square deviation from 0 and were used for further analysis.

**Figure S3.**
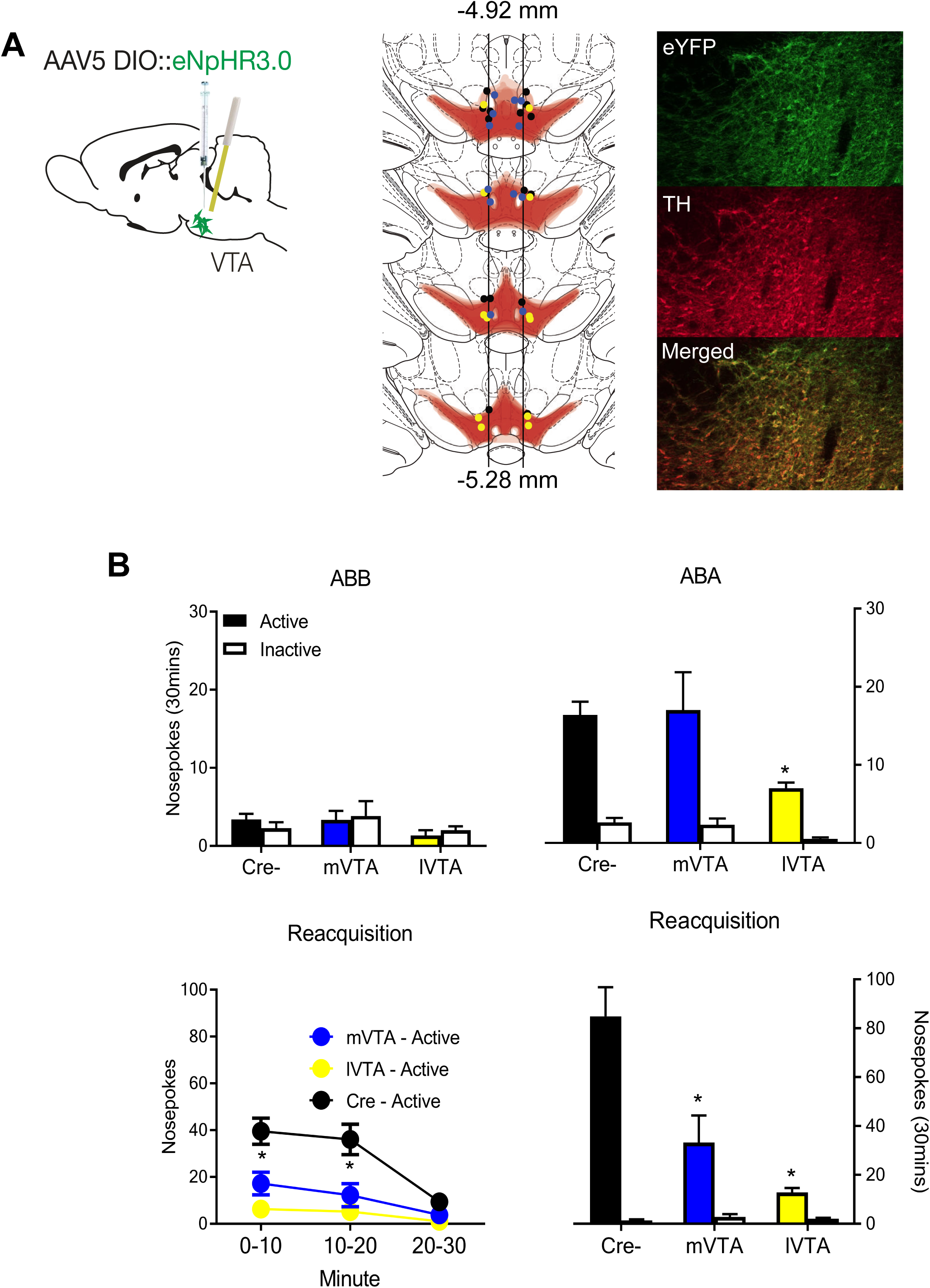
VTA optogenetic inhibition and relapse. (**A**) Location of eNpHR3.0 expression and fibre tips in midbrain with each rat represented at 25% opacity. The lateral edge of the fasciculus retroflexus was used as anatomical boundary between mVTA and lVTA. Example of eNpHR3.0 expression in VTA TH neurons. (**B**) Mean ± SEM nosepokes during test in extinction and training contexts. Rats refrained from alcohol-seeking in the extinction context but relapsed to alcohol-seeking in the training context (context main effect: F (1, 17) = 33.83, p < 0.001; nosepoke main effect F (1, 17) = 32.68, p < 0.001; context x nosepoke interaction, F (1, 17) = 43.30, p < 0.001) (**Figure 1B**), showing renewal (context-induced reinstatement) of alcohol-seeking. lVTA optogenetic inhibition around Nosepoke+ reduced responding on ABA test (main effect group F (1, 17) = 8.93, p = 0.008; group x context interaction F (1, 17) = 5.11, p = 0.037) whereas mVTA inhibition did not (main effect group F (1, 17) = 0.17, p = 0.69; group x context interaction F (1, 17) = 0.09, p = 0.77). Mean ± SEM nosepokes during reacquisition test. lVTA (main effect group F (1, 17) = 25.18, p < 0.001; group x nosepoke interaction F (1, 17) = 26.22, p < 0.001) and mVTA (main effect group F (1, 17) = 12.47, p = 0.003; group x nosepoke interaction F (1, 17) = 14.04, p = 0.002) inhibition around Nosepoke+ reduced reacquisition. * p < .05.

**Figure S4.**
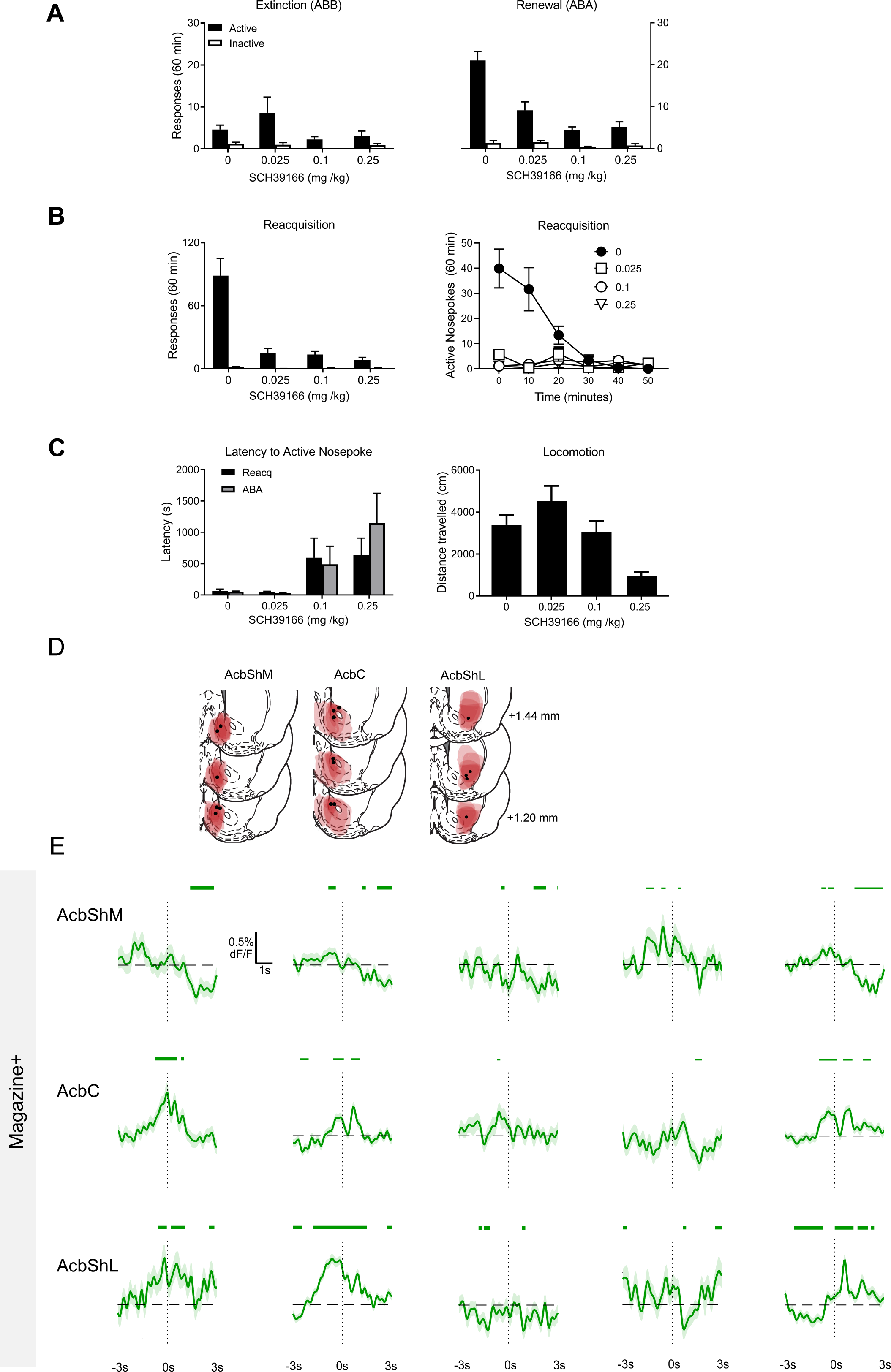
Dopamine and relapse. (**A**). **Effects of SCH39166 on renewal**. Mean ± SEM nosepokes during test in extinction and training contexts. Rats refrained from alcohol-seeking in the extinction context but relapsed to alcohol-seeking in the training context (context main effect: F (1, 28) = 21.85, p < 0.001; nosepoke main effect F (1, 28) = 51.55, p < 0.001; context x nosepoke interaction, F (1, 28) = 27.56, p < 0.001). SCH39166 reduced renewal (context x group interaction: F (1, 28) = 29.13 and context x nosepoke x group F (1, 28) = 44.86, p < 0.001), including at the lowest dose (0.025 m/kg) (context x group interaction: F (1, 28) = 21.69, p < 0.001 and context x nosepoke x group F (1, 28) = 35.49, p < 0.001). (**B**). **Effects of SCH39166 on reacquisition.** Mean ± SEM nosepokes during test for reacquisition. SCH39166 reduced reacquisition (group main effect: F (1, 28) = 57.40 and group x nosepoke interaction F (1, 28) = 59.66, p < 0.001), including at the lowest dose (0.025 m/kg) (0 v 0.025 mg/kg main effect: F (1, 28) = 35.87, p < 0.001 and group x nosepoke F (1, 28) = 36.46, p < 0.001). (**C**). **Effect of SCH39166 on latency to first active nosepoke and locomotor activity**. There was an overall effect of SCH39166 on latency to first active nosepoke (main effect of drug F (1, 28) = 4.55, p = 0.04) and a linear effect of SCH39166 dose (F (1, 28) = 11.72, p = 0.0019). Importantly, there was no effect of the lowest dose of SCH39166 (F (1, 28) = 0.006, p = 0.939) even though this dose reduced relapse. There were no interactions with test session (ABA, Reacquisition). There was no overall effect of SCH39166 on locomotor activity (main effect F (1, 28) = 0.84, p = 0.37). Locomotor activity was reduced at the highest dose (0.25 mg/kg) (linear trend across dose: F (1, 28) = 23.91, p < 0.001). Critically, the lowest dose (0.025 mg/kg) which reduced relapse did not affect locomotor activity compared to 0 mg/kg (F (1, 28) = 2.43, p = 0.13). (**D**)**. dLight Histology**. Location of dLight1.1 expression and fibre tips in nucleus accumbens with each rat represented at 25% opacity (**E**). **dLight transients**. Mean ± SEM dopamine transients from -3s to 3 s around Magazine+ for AcbShM, AcbC, AcbShL. Green bars above traces show periods (with minimum consecutive threshold) of significant difference from 0 as defined by 95% confidence intervals.

